# ShinyCell2: An extended library for simple and sharable visualisation of spatial, peak-based and multi-omic single-cell data

**DOI:** 10.1101/2025.04.22.650045

**Authors:** Bei Jun Chen, Yi Yang Lim, Xinyi Yang, Lijin Wang, Owen J L Rackham, John F Ouyang

**Author notes:** Correspondence to (OJLR) and (JFO).

## Abstract

Single-cell technologies now span multiple modalities, generating large, complex datasets that challenge analysis and sharing. We present **ShinyCell2**, an enhanced R package for interactive visualisation of single-cell multi-omics and spatial transcriptomics data. **ShinyCell2** retains the simplicity and lightweight deployment of its predecessor while introducing advanced visualisations, cross-modality comparisons, and statistical tools tailored to spatial and multi-omic data. It enables intuitive, rapid exploration of high-dimensional data without requiring extensive computational expertise.

## Main

Single-cell datasets are inherently complex and typically require bioinformatics expertise to query and visualise gene expression patterns. Moreover, these datasets are often massive in size, making them challenging to share and distribute efficiently. This problem is exacerbated by the increasing complexity and requirements of analysis tools. To address these limitations, there is a growing need to deploy large single-cell datasets as online interactive applications. Such platforms enable biologists to intuitively explore and analyse the data in real time, without the burden of downloading and processing the full dataset locally. To this end, we previously developed ShinyCell,^1^ a lightweight R package that generates explorable and shareable interactive web interfaces from single-cell RNA sequencing data. ShinyCell enables intuitive data exploration, particularly for researchers without extensive computational expertise, and supports seamless deployment for both local and web-based platforms.

In recent years, single-cell technologies have advanced to encompass multiple modalities, such as CITE-seq^2^ for simultaneous profiling of RNA and protein expression, scATAC-seq^3^ for mapping open chromatin regions, and scCUT&Tag^4^ for profiling histone modifications. Also, spatial transcriptomics now enables the study of the spatial organisation of cell types within complex tissues. Consequently, existing online interactive applications must evolve to support these emerging data types, each of which requires specialised visualisations. In response to this need, we present ShinyCell2, a major extension of our original platform that broadens its capabilities to accommodate multi-modal single-cell and spatial datasets. The software is publicly available at https://github.com/the-ouyang-lab/ShinyCell2.

The ethos underpinning ShinyCell—easy generation from major data formats, lightweight and portable deployment, and customisable aesthetics—is preserved and extended. ShinyCell2 integrates seamlessly within the existing landscape of analysis tools such as Seurat,^5^ ScanPy,^6^ ArchR,^7^ and Signac^8^ (Figure 1a). It uniquely targets users seeking straightforward, customisable exploration of their datasets without the complexity of extensive setup or external dependencies. Unlike alternative tools that require complex configuration,^9–11^ ShinyCell2 allows straightforward local deployment and is tightly integrated with R/Bioconductor workflows (Figure 1b).

**Figure 1:**
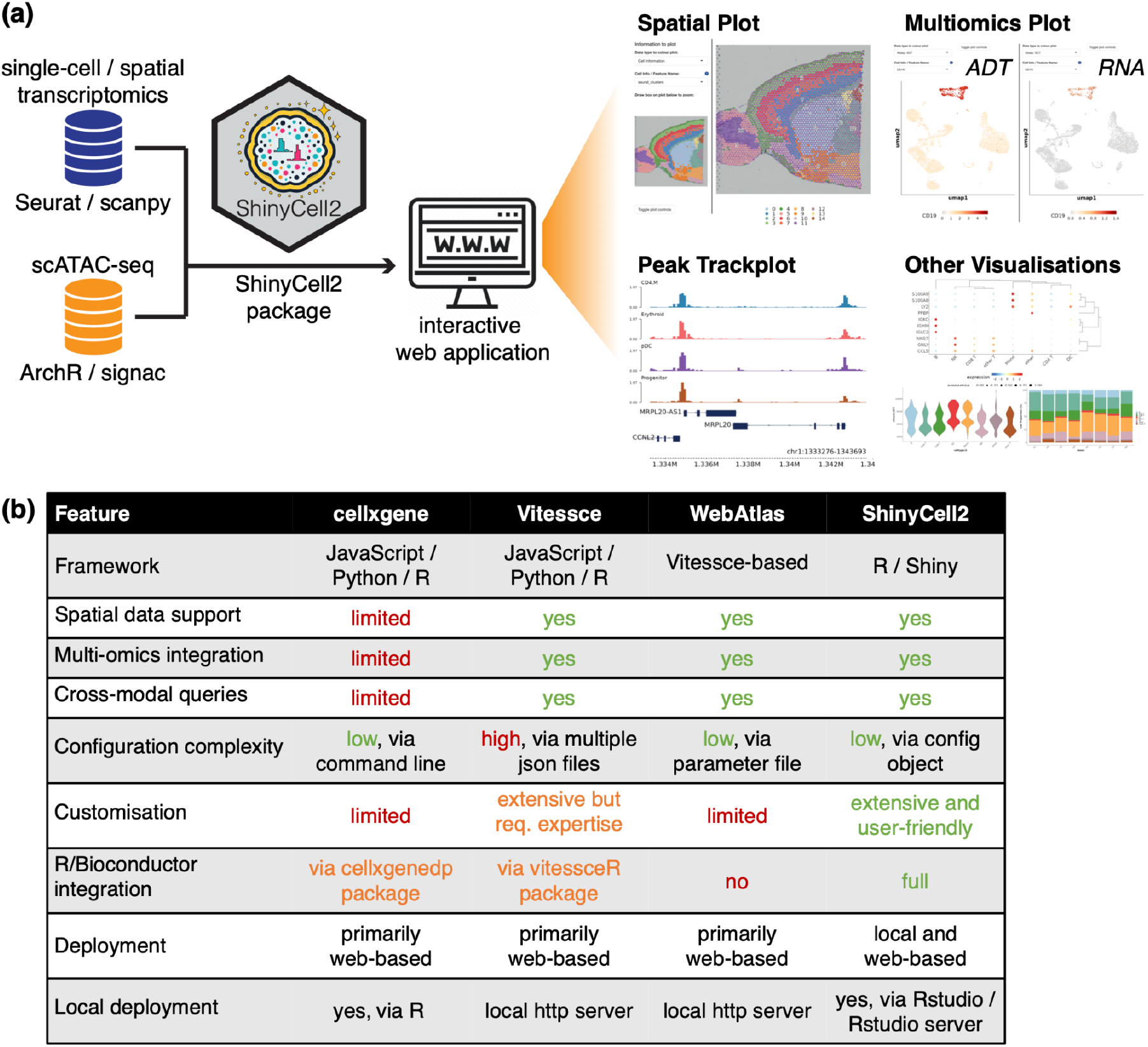
(a) Schematic showing the ShinyCell2 workflow involving the generation of interactive web applications from common single-cell / spatial data formats and example visualisations in the web applications. (b) Comparison of key features in ShinyCell2 with other existing single-cell web application generation tools.

ShinyCell’s approach to visualisation emphasises the parallel exploration of observations e.g. gene expression or peak accessibility and cell-level metadata, allowing users to explore and uncover patterns in their data. ShinyCell2 enriches this concept with enhanced visualisation capabilities. New features include spatial plots to elucidate gene expression within tissue architecture (Figure 2a), and peak track plots for exploring epigenetic landscapes from scATAC-seq data (Figure 2b). Furthermore, side-by-side UMAPs and scatter plots facilitate cross-modality analyses (Figure 2c–d), such as comparing RNA expression and protein levels. Additionally, metadata confusion matrices aid in elucidating hierarchical relationships within cell-type annotations, making it easier to identify subtle cellular subpopulations (Figure 2e). The visualisations available in the original ShinyCell, such as cell proportion plots, violin plots, and gene expression bubbleplots, are retained in ShinyCell2 (Figure 2f–h).

**Figure 2:**
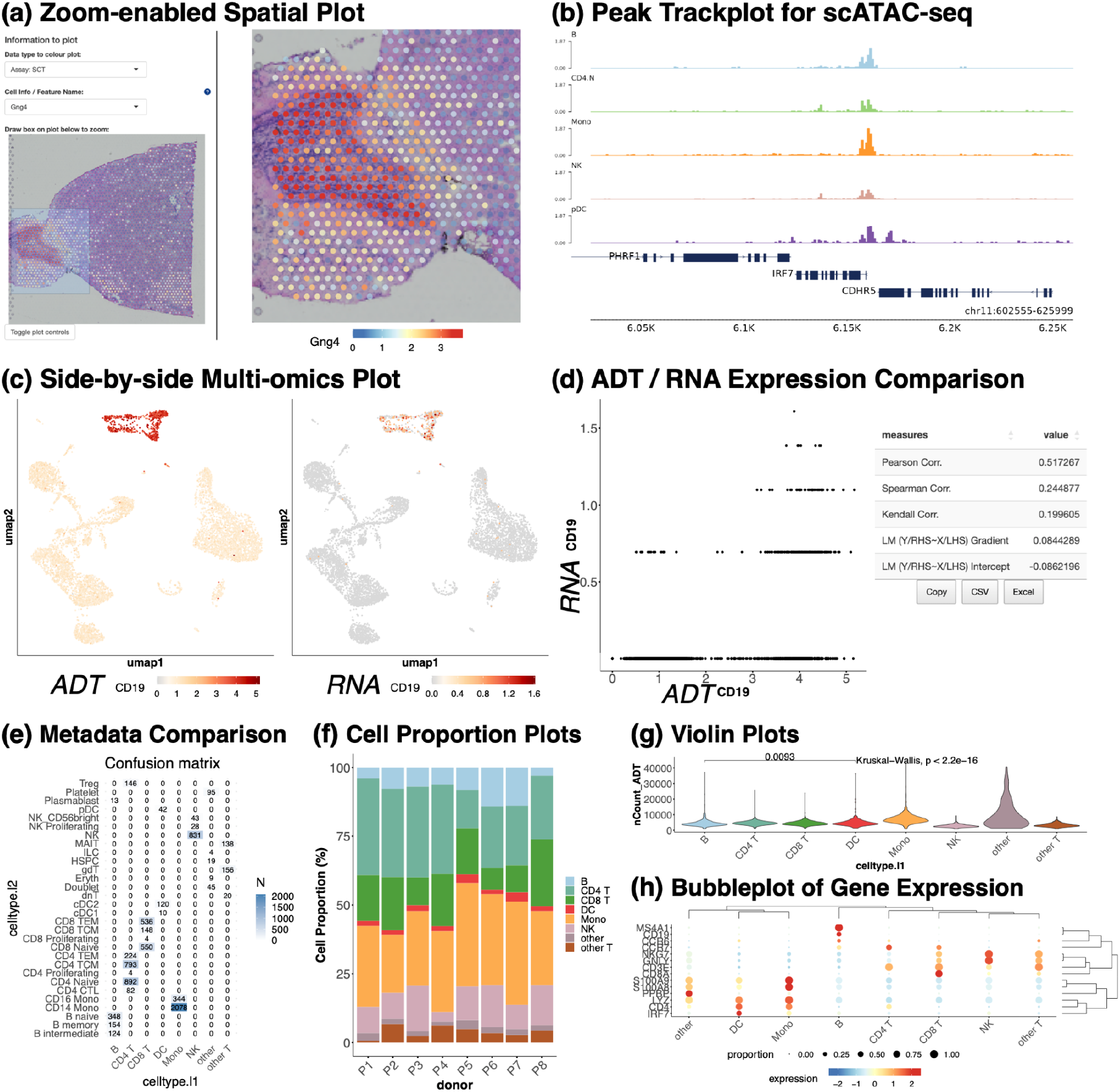
Examples of new visualisation options in ShinyCell2 including (a) zoom-enabled spatial plots, (b) peak trackplot, (c,d) side-by-side multi-omics dimension reductions and comparisons and (e) comparison of cell metadata via confusion matrix visualisation. Plots available in the original ShinyCell are retained in ShinyCell2 including (f) cell proportion plot, (g) violin plot, and (h) gene expression bubbleplot. Note that all the images in this figure are taken directly from a showcase ShinyCell app https://shinycell.ouyanglab.com/, illustrating the utility of these interactive web applications.

ShinyCell2 also introduces substantial functional enhancements. These include dynamic, zoomable UMAP visualisations, allowing detailed inspection of cell clusters without loss of resolution and side-by-side plotting from multiple assays (e.g., RNA vs. protein), facilitating intuitive cross-assay comparisons. Advanced data representation includes percentile-based min / max cutoffs to manage outliers, improving clarity in visualisation effectively. Improved figure management ensures consistent plot sizing, supporting direct comparisons and the generation of publication-ready figures. Furthermore, built-in statistical testing, such as global ANOVA or pairwise comparisons, enable rigorous quantitative analyses between cell populations. The flexibility of data visualisation is also enhanced through options to reorder summary data, such as cluster proportion plots, assisting in quickly identifying dominant cell types or conditions across different samples. Finally, comprehensive customisation options for colours, scales, sizes and text labels throughout allow researchers to reproduce the visualisation closely aligned with those in their publications.

ShinyCell2 can be applied to a wide range of experimental designs, providing its capabilities in integrative analyses across transcriptomic, epigenomic and spatial data. For example, it can effectively visualise which cell-type-specific regulatory elements (using scATAC-seq data) are linked with gene expression programs (from single-cell RNA sequencing). Equally, ShinyCell2 can overlay gene expression with protein abundance (from CITE-seq) and spatial location (using spatial transcriptomics), facilitating direct exploration of RNA-protein relationships at the spatial level. Through these enhanced visualisation and integrative functionalities, ShinyCell2 empowers researchers to dissect complex biological systems. The tool provides robust visualisations for examining tumour microenvironments, understanding cellular differentiation pathways, and exploring cellular heterogeneity with high resolution.

These visualisations are enabled by a lightweight data storage and retrieval system built on the HDF5 format. Relevant observations and metadata are extracted from analysis-specific file formats e.g. Seurat objects, indexed, and stored on disk for efficient retrieval. This design eliminates the dependency on the original analysis packages, allowing the resulting web application to remain lightweight. As a result, even large datasets can be visualised and shared without requiring memory-intensive computing environments.

In conclusion, ShinyCell2 represents a comprehensive upgrade aligning with contemporary single-cell methodologies, extending functionality to spatial transcriptomics, CITE-seq, and scATAC-seq data. True to the ShinyCell philosophy, it provides powerful visualisation features in a user-friendly, easily deployable interface while maintaining a low computational footprint and supporting seamless customisation and extension. ShinyCell2 thus significantly enhances researchers’ ability to interpret and share complex single-cell multi-omics and spatial datasets, driving new insights in cellular biology and disease mechanisms.

## Online Methods

### ShinyCell2 Overview

ShinyCell2 is an R package that builds upon the foundation of ShinyCell^1^ to enable interactive visualisation and analysis of complex single-cell data via Shiny-based web applications. ShinyCell2 extends the functionality of its predecessor by supporting the simultaneous exploration of diverse data modalities, including spatial transcriptomics, CITE-seq, scATAC-seq, and conventional scRNA-seq. The package is optimised for large-scale datasets by maintaining a low memory footprint. Specifically, it leverages the hdf5r R package to read only the necessary data e.g. user-specified cell metadata and assay expression from HDF5 files instead of loading the entire single-cell data object into memory.

### ShinyCell2 App Generation Workflow and Supported Filetypes

ShinyCell2 applications can be generated with only five lines of code as follows:

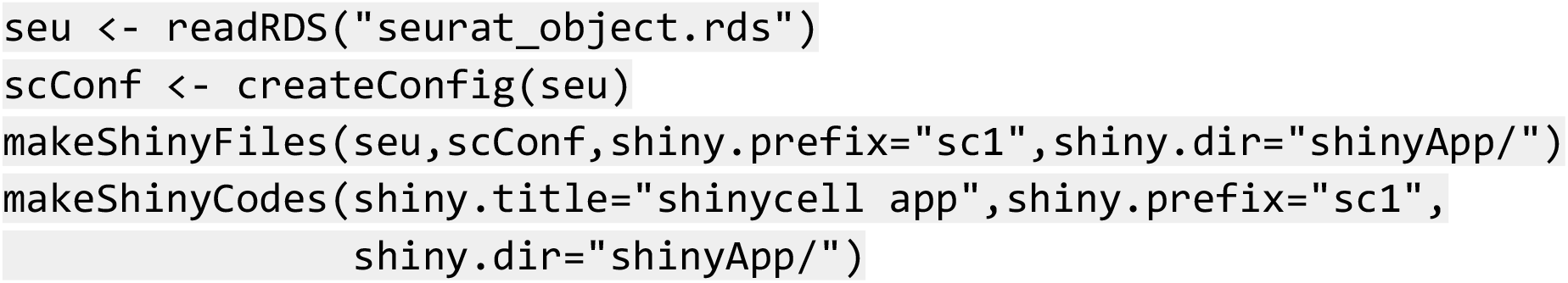

The first two lines of code load the processed single-cell data object, typically a Seurat object in this case. Next, a ShinyCell2-specific configuration object is generated using `createConfig()`, which stores metadata display settings such as labels and colour palettes. The ShinyCell2 configuration and single-cell object are then used to generate the files and code required for the ShinyCell2 app via the `makeShinyFiles()` and `makeShinyCodes()` functions respectively. The shiny.prefix argument is used to define a unique file prefix for each dataset, and this prefix must be consistent across both functions. This mechanism allows for the integration of multiple single-cell or spatial datasets into a single ShinyCell2 app. To do so, `makeShinyFiles()` is run separately on each dataset with a distinct `shiny.prefix` and `makeShinyCodes` is run once to combine all the datasets into a unified app. The code below demonstrates how to create a multi-dataset ShinyCell2 app with two datasets:

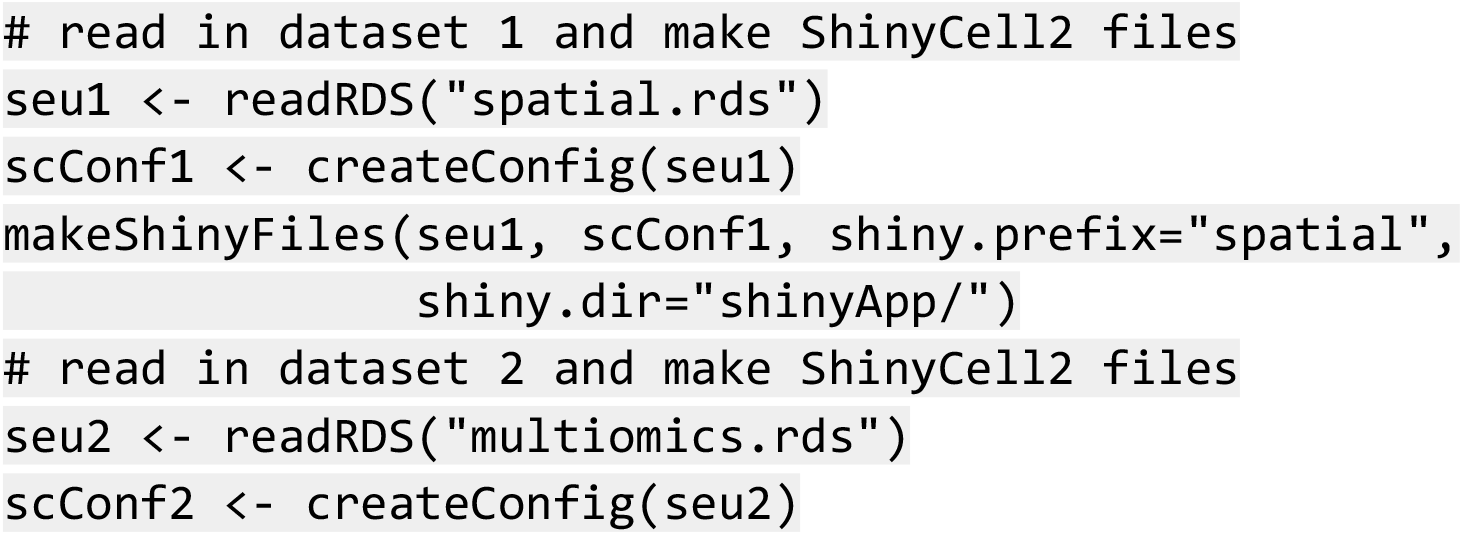
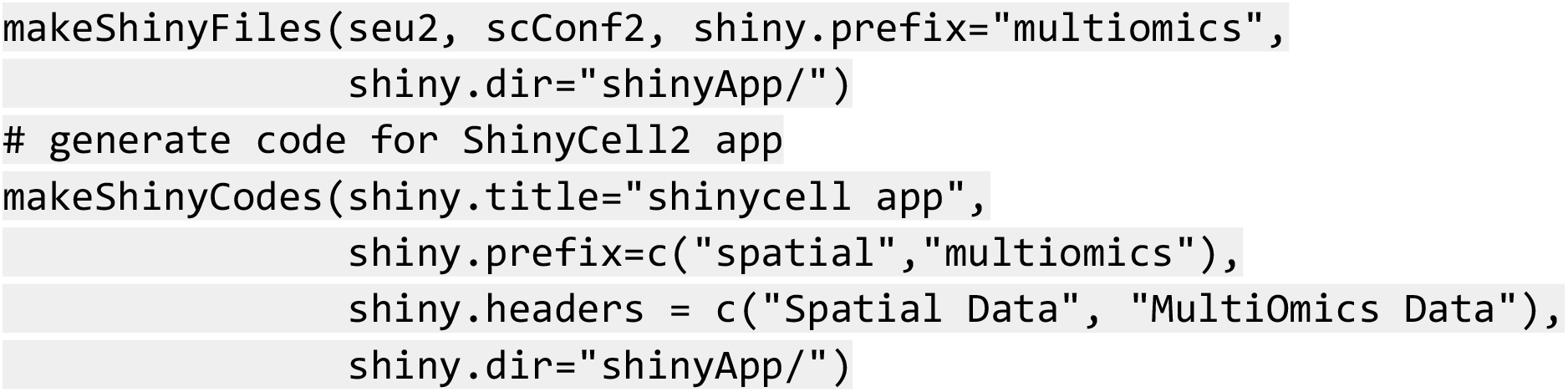

Building on the original ShinyCell, the app’s appearance and content can be tailored using the ShinyCell2 configuration. Users can selectively remove metadata fields or rearrange their display order to streamline and declutter the metadata. Labels and colour palettes for categorical metadata e.g. cluster identities can be customised to match the visual style used in publication figures. By default, ShinyCell automatically selects a set of metadata and genes for visualisation, but these defaults are easily modifiable, giving users full control over the app’s initial display settings.

In terms of input data, ShinyCell2 supports common single-cell RNA-sequencing, multi-omics and spatial data formats, including R-based objects from the Seurat,^2^ Signac,^3^ and ArchR^4^ packages, as well as h5ad files from the Python-based scanpy toolkit.^5^ Other formats e.g. SingleCellExperiment or loom can be readily converted to compatible formats (e.g., Seurat or h5ad) using the sceasy R package, allowing them to be seamlessly integrated into the ShinyCell2 app generation workflow.

### ShinyCell2 App File Structure

The ShinyCell2 app consists of two main components: (i) memory-optimised files containing single-cell metadata and assay expression data, and (ii) a set of R scripts that power the interactive visualisations within the Shiny interface. To enhance performance, single-cell metadata is stored in data.table format, which offers faster data access compared to the default data.frame. Assay expression matrices are saved in HDF5 format, enabling on-demand loading of only the genes or features specified by the user. This avoids loading the entire dataset into memory, significantly reducing memory usage and improving the responsiveness of the app. For peak track visualisations, bigWig files are used to store peak coverage information. Widely used in genome browsers, the bigWig format is compact and supports efficient, region-specific access without requiring the full file to be loaded into memory. All these optimisations operate on the backend, providing a seamless user experience. Moreover, the code is made transparent to users, providing them with the flexibility to extend the app by incorporating custom visualisations or analysis modules tailored to their specific single-cell workflows.

### Spatial scRNA-seq Visualisation

Spatial scRNA-seq data can be visualised through the “Zoom-enabled Spatial” and “Side-by-side Spatial” tabs. The Zoom-enabled Spatial tab allows users to overlay cell metadata or gene expression data onto tissue images, with zoom functionality for detailed exploration of specific regions. The zooming is achieved by restricting the viewpoint of the underlying ggplot2 object. This approach ensures quick response with no loss of resolution, allowing rapid and detailed exploration of specific regions. The Side-by-side Spatial tab enables direct comparison of cell metadata e.g. celltype annotations, with gene expression within the spatial context of the tissue architecture.

### scATAC-seq Track Plot Visualisation

For scATAC-seq data, ShinyCell2 incorporates a previously published Trackplot function^6^ to generate IGV-style locus plots for visualising open chromatin regions. The Trackplot function is designed to be lightweight and efficient, relying only on the data.table R package and bwtool,^7^ a command-line utility for querying bigWig files. By leveraging bwtool, the function enables rapid, region-specific extraction of accessibility signals directly from bigWig files, achieving performance that is over 15 times faster than comparable plotting functions. This speed advantage is particularly beneficial when working with large datasets or rendering multiple regions in a single session. Users can specify regions of interest by gene name or genomic coordinates, and enhance the plots with custom annotation tracks, such as candidate cis-regulatory elements (CREs) from ENCODE,^8^ provided in standard BED format. This flexibility allows for rich, context-aware visualisations that integrate accessibility signals with known regulatory features.

### Multi-Assay Visualisation and Integration

ShinyCell2 supports multi-assay single-cell data visualisation and integration, enabling seamless switching between molecular layers such as RNA expression, protein abundance, and chromatin accessibility. This facilitates the exploration of regulatory mechanisms and cellular heterogeneity within a unified interface. The app provides interactive UMAP visualisations of multi-omics profiles, flexible feature selection across assays, and tools for comparing cell populations across different data types. For example, users can assess the consistency of cell type annotations between RNA, protein, and chromatin-based classifications. Together, these features offer a comprehensive and integrated view of single-cell biology by enabling the simultaneous analysis of multiple molecular features within individual cells.

### Enhanced Visualisation Features

ShinyCell2 introduces a slew of enhancements to support advanced data visualisation and analysis in single-cell studies. First, the Zoom-enabled UMAP feature allows users to zoom into specific regions of UMAP plots without any loss of resolution. This enables simultaneous global and local perspectives, aided by dynamic displays of cell counts in both the full and zoomed-in views. Second, to improve the clarity and usability of visual outputs, plot layouts have been enhanced, where legends are now rendered separately from the main plots. This ensures consistent sizing for side-by-side visualisations and allows users to export the main plots and legends as separate PNG or PDF files for flexible post-processing. Third, data points with extreme values can distort the colour scale by inflating its range, making it difficult to interpret the visualisation. To address this, ShinyCell2 applies percentile-based minimum and maximum cutoffs, effectively minimising the influence of outliers and enhancing the clarity of colour-based data representations. Fourth, in the “Side-by-side DimRed” tab, users can perform comparative visualisations across different cell metadata and assay expressions. This includes confusion matrices to compare cell memberships between categorical variable pairs or scatter plots with correlation statistics for continuous variables. Fifth, the “Violin / Boxplot” tab integrates statistical testing, supporting both global and pairwise comparisons across groups, enabling users to draw robust analytical conclusions directly within the app. Sixth, the “Proportion plot” tab includes flexible ordering of the X-axis, allowing users to sort groups by cell proportions. This makes it easier to identify clusters that are dominated by cells from specific biological or experimental conditions. Finally, the “Bubbleplot / Heatmap” tab offers customizable colour scales through max value clipping, allowing users to adjust the dynamic range of the visualisation and better highlight relevant data patterns.

## Code and Data Availability

The ShinyCell2 R package is available publicly on Github at https://github.com/the-ouyang-lab/ShinyCell2. Comprehensive tutorials covering app creation, aesthetic customisation, and web deployment are provided on the GitHub repository. An example web app showcasing the visualisation of spatial transcriptomics, scATAC-seq, and CITE-seq datasets can be accessed at https://shinycell.ouyanglab.com/, allowing users to explore the full functionality of ShinyCell2. The datasets used in the example app are available on Zenodo at https://zenodo.org/records/15162323.

